# Whole-body single-cell sequencing of the *Platynereis* larva reveals a subdivision into apical versus non-apical tissues

**DOI:** 10.1101/167742

**Authors:** Kaia Achim, Nils Eling, Hernando Martinez Vergara, Paola Yanina Bertucci, Thibaut Brunet, Paul Collier, Vladimir Benes, John C Marioni, Detlev Arendt

## Abstract

Animal bodies comprise a diverse array of tissues and cells. To characterise cellular identities across an entire body, we have compared the transcriptomes of single cells randomly picked from dissociated whole larvae of the marine annelid Platynereis dumerilii^1–4^. We identify five transcriptionally distinct groups of differentiated cells that are spatially coherent, as revealed by spatial mapping^5^. Besides somatic musculature, ciliary bands and midgut, we find a group of cells located at the apical tip of the animal, comprising sensory-peptidergic neurons, and another group composed of non-apical neural and epidermal cells covering the rest of the body. These data establish a basic subdivision of the larval body surface into molecularly defined apical versus non-apical tissues, and support the evolutionary conservation of the apical nervous system as a distinct part of the bilaterian brain^6^.

## Introduction

Animal bodies are composed of cells, which are organized into tissues and organs. Recently, a number of studies have applied single cell transcriptomics to assess cellular diversity within tissues, such as the vertebrate pancreas^7^, intestine^8^, or different brain parts such as cortex, basal ganglia or hypothalamus^9–11^. This approach allows the molecular characterisation of cell types within a given tissue, as well as an assessment of their heterogeneity. However, the sheer number of cells in conventional vertebrate or insect systems has so far hampered the comparison of cellular identities across tissue boundaries, or even across entire bodies.

Other laboratories have focused on the molecular comparison of different tissues, referred to as tissue transcriptomics. This approach determines and compares the expression profiles of entire tissues via bulk RNAseq^12,13,14,15^, with the potential to compare across the entire body. Inherent to the approach however, tissue transcriptomics so far refer to the somewhat arbitrary, morphology-based dissection of the body into discernible units. This is relevant because, on the one hand, cell types of a given tissue or organ may be very divergent and on the other hand, similar cell types can populate different tissues. Therefore, while tissue transcriptomics allow in-depth and technically robust analyses and comparisons of larger cellular assemblies, these are somewhat arbitrarily defined and the method lacks the resolution to disentangle and molecularly define cell and tissue identities across entire bodies.

To advance on this, we establish the marine annelid *Platynereis dumerilii*, a molecular model species for development, evolution and neurobiology^16,17^, as experimental paradigm to explore how cellular identities compare and relate to each other across an entire animal body. We apply single-cell RNAseq to randomly sampled cells from the dissociated whole larvae at 48 hours post fertilization (hpf). The *Platynereis* larval body is especially suited for this task, as it already comprises various cell types and tissues. Besides larval body features (e.g., epidermal cells and ciliary bands) it contains early differentiating cells contributing to the later juvenile worm (musculature, brain and nerve cord)^18^. It thus allows us to determine and compare cellular identities at an early stage of the life cycle when cell numbers are still relatively low.

Our whole-body analysis reveals that, at this stage, the larval annelid body comprises five groups of differentiated cells with distinctive expression profiles. In each of these groups, cells express sets of transcripts that together encode group-specific cellular modules – representing cellular structures and functions that characterize these cells. While some of these groups and their defining modules, such as ciliary bands with motile cilia or larval musculature with striated myofibres, match larval morphology, others shed new light on the basic organization of the annelid body. We find a group of cells located at the apical tip of the animal that appears to be distinct from all other ectodermally-derived, non-apical cells of the body. While the apical cells specifically utilize photosensory and neurosecretory modules, the non-apical tissue instead shows specific expression of various extracellular matrix components – such as ligand-receptor pairs, cellular adhesion molecules, and conserved proteolytic enzymes. Comparative data indicate that this molecular subdivision into apical versus non-apical tissues is an important organisational feature of the body that appears to be conserved in animal evolution.

## Results

### Single-cell RNA-seq identifies five groups of differentiated cells

To explore cell type diversity on the whole organism level, we dissociated whole larvae of a marine annelid, *P.dumerilii* at 48 hpf and randomly captured cells for single-cell RNA-sequencing (scRNA-seq) (**Fig. 1**). At this stage of development, the larva comprises of relatively few cells (~5000), but many differentiated cell types such as ciliated cells, neurons or myocytes are already present. The collected cells were optically inspected to exclude doublets, multiple cells or cell debris. Sequenced samples were further filtered computationally to remove low complexity transcriptomes, lowly-expressed genes and transcriptomic doublets (**Extended Data Fig. 1**, Methods). A total of 373 cells and 31300 transcripts passed filtering steps and were used for downstream analysis. To group the cells into distinct clusters we used a sparse clustering strategy, which identified seven groups of cells. We used the *scran* package to find group specific marker genes and discovered that in pairwise comparisons across all groups, two clusters were consistently highly similar to one other. Therefore, we merge these two closely related groups (**Fig. 1, Extended Data Fig. 2**, Methods).

**Figure 1.**
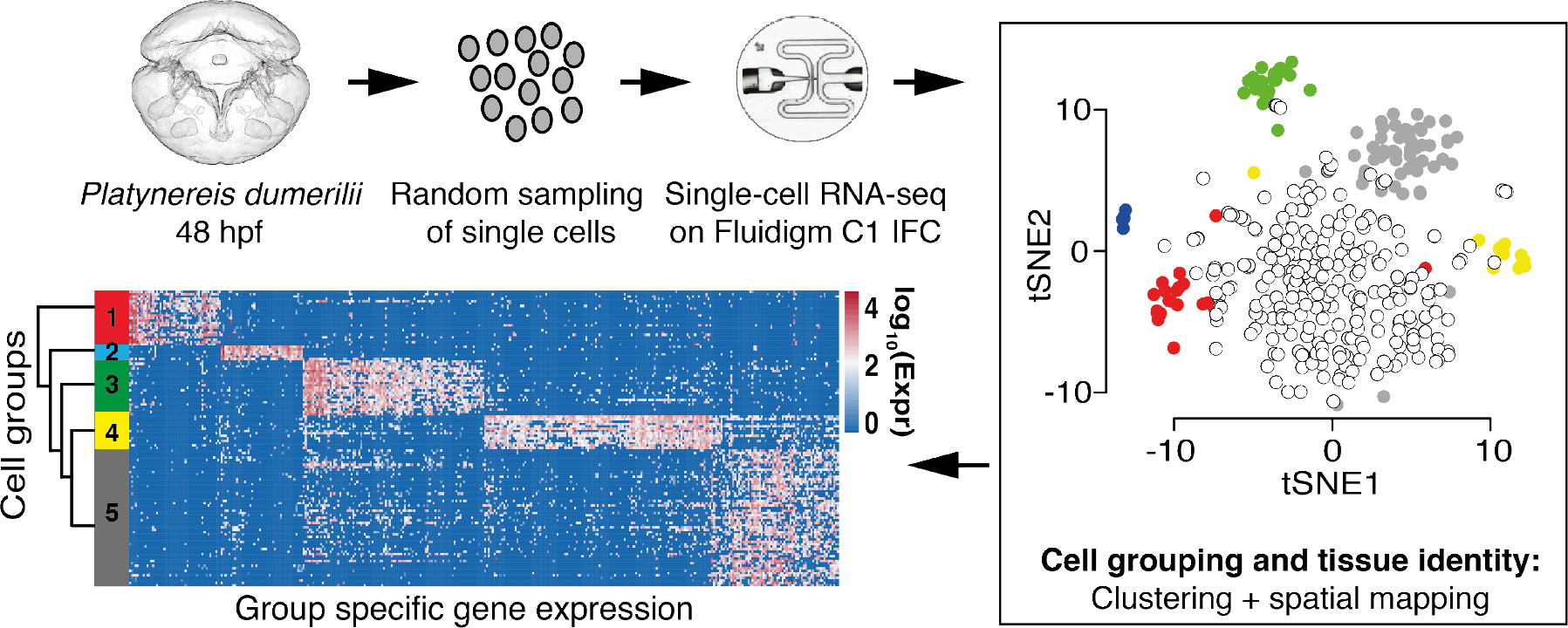
Single cell transcriptomics of *Platynereis* 48 hpf larvae. Cells of the 48 hpf larvae were dissociated and randomly selected for single cell RNA-sequencing using the Fluidigm C1 Single-cell AutoPrep system. Combining sparse clustering with spatial positioning of single cells allows the identification of robust cell groups within the data. The clustering approach enables identification of genes that characterise each cell type. Finally, we used hierarchical clustering to investigate the similarity between the identified cell clusters.

To characterize the remaining six groups further, we identified differentially expressed genes (Methods). The largest group of cells was characterized by the specific expression of genes known to be active in developmental precursors, such as DNA replication (*DNA topoisomerase, Rfc3, Rfc5, Mcm3, Mcm4, Mcm7*), cell proliferation (*Pcna, Md2l1, Apc7*, **Extended Data Fig. 2i**), or chromatin remodeling (*Nucleoplasmin, Bptf*). We thus inferred that this group was comprised of undifferentiated cells.

Cells in the other five groups showed significantly lower expression of these markers (**Extended Data Table 1**, FDR < 0.1, Methods) and we thus consider them differentiated cells. For each of these five groups, pairwise differential expression analysis revealed distinct sets of group-specific effector genes (i.e., differentially expressed genes that encode the particular structural and/or functional properties of cells) and group-specific transcription factors, many of which are known to act as terminal selectors in cell type differentiation^19,20^ (**Extended Data Table 2**).

To validate the clustering of differentiated cells into the above groups, and to infer relationships between these groups, we used bootstrap resampling (Methods) to calculate a hierarchical tree (**Extended Data Fig. 3**). This tree confirmed each group’s integrity, albeit with variable support. We then averaged the gene expression across cells in each group and used these values to calculate a tree comprising cell groups only (**Fig. 3**, upper panel). The resulting tree retrieved the group topology of the single cell tree with higher support values.

### Spatially coherent, molecularly defined larval tissues

To characterize and localize the five distinct groups of differentiated cells with regard to larval morphology, we used a dual strategy. First, we investigated the spatial distribution in the larval body by mapping constituent cells into a cellular-resolution expression atlas^5^ (**Fig. 2**). To this end, we constructed a cellular resolution atlas of the 48 hpf larva (**Extended Data Fig. 4**), taking advantage of the existing Profiling by Signal Probability mapping (ProSPr) pipeline^21^. Using ProSPr, the majority (95%) of cells could be mapped to distinct regions within the larva. Complementing this, we used wholemount in situ hybridization (WMISH) with probes targeted towards group-specific transcripts to refine the spatial mapping.

**Figure 2.**
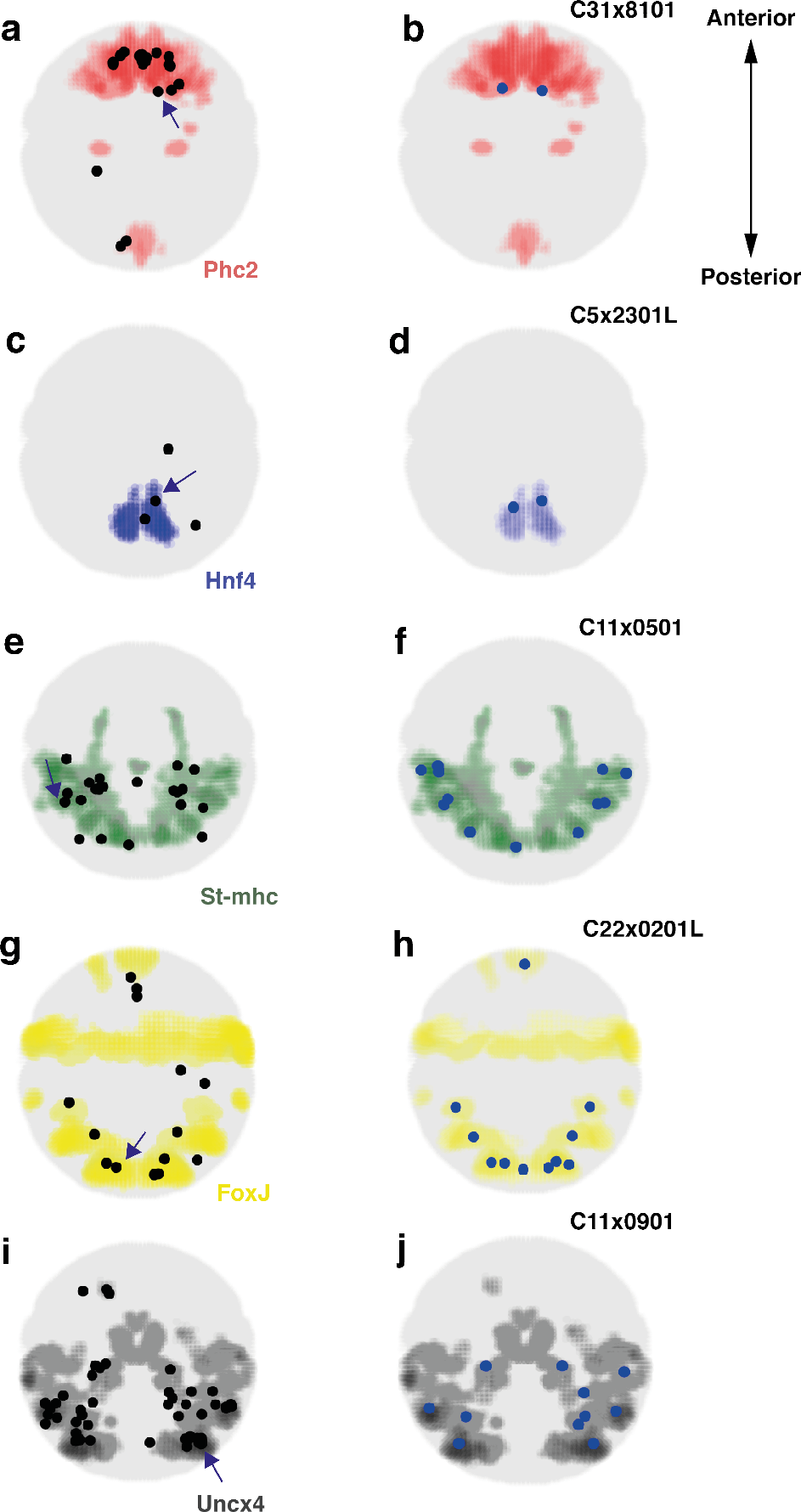
Spatial mapping of single cells to tissues characterized by specific marker gene expression. **a-j**, Left panels: Centroids of the voxel clusters to which cells were mapped with highest confidence are shown as black dots. Tissue specific marker gene expression is visualized as colored regions within the embryo. Left panels: Mapping results for all cells belonging to the respective group. Right panels: For each cell group, we show an example of the mapping result for one individual cell (indicated by an arrow on the left panel). **a**, Cells of the apical ectoderm group (n = 23 cells) map to the *Phc2* expressing region in the embryo. **b**, Mapping result for the cell C31x8101. **c**, Midgut cells (n = 4 cells) map to the *Hnf4* expressing region. **d**, Mapping result for the cell C5x2301L. **e**, Striated musculature cells (n = 23 cells) map to the *St-mhc* expressing region. **f**, Mapping result for the cell C11x0501. **g**, Cells of the ciliary bands (n = 14 cells) map to the *Foxj* expressing region. **h**, Mapping result for the cell C22x0201L. **i**, Cells of the non-apical ectodermal group (n = 55 cells) map to the *Uncx4* expression domain. **j**, Mapping result for the cell C11x0901.

Following this strategy, cells from group 1 mapped to the most apical part of the larval ectoderm around the so-called apical organ (**Fig. 2a-b**). Consistent with this, the specific group 1 marker *Phosphodiesterase-9* (*Pde9*) outlined a highly specific expression territory in the apical/dorsal episphere (**Fig. 3a**). With reference to the clear neural character of the cells in this group (see below), we refer to group 1 cells as ‘apical nervous system’. Group 2 cells mapped into the yolk-rich macromeres where the first differentiating cells of the later midgut are located (**Fig. 2c-d**), as confirmed by the highly specific expression of *Hnf4* in the cellularized portion of the macromeres (**Fig. 3b**). This group thus represents ‘larval midgut’. Cells of group 3 mapped to the differentiating striated myocytes (**Fig. 2e-f**) and, in line with this, *Striated muscle myosin heavy chain (St-mhc)* was expressed in all cells belonging to ‘striated musculature’ (**Fig. 3c**). Next, group 4 cells mapped to the ciliary bands (**Fig. 2g-h**), which is composed of multiciliated cells with motile cilia. Expression of *Radial spoke head protein homolog 4 (Rsph4)* in the multiciliated cells of the ciliary bands of the apical organ, prototroch and paratrochs confirmed this mapping (**Fig. 3d**)^18^. Complementing this, group 5 cells covered much of the remaining larval surface, including non-apical territories in the larval trunk (**Fig. 2i-j**). The broad expression of the *Metabotropic glutamate receptor 7 (Grm7)* in trunk and head clearly illustrates these ‘non-apical surface cells’ (**Fig. 3e**).

**Figure 3.**
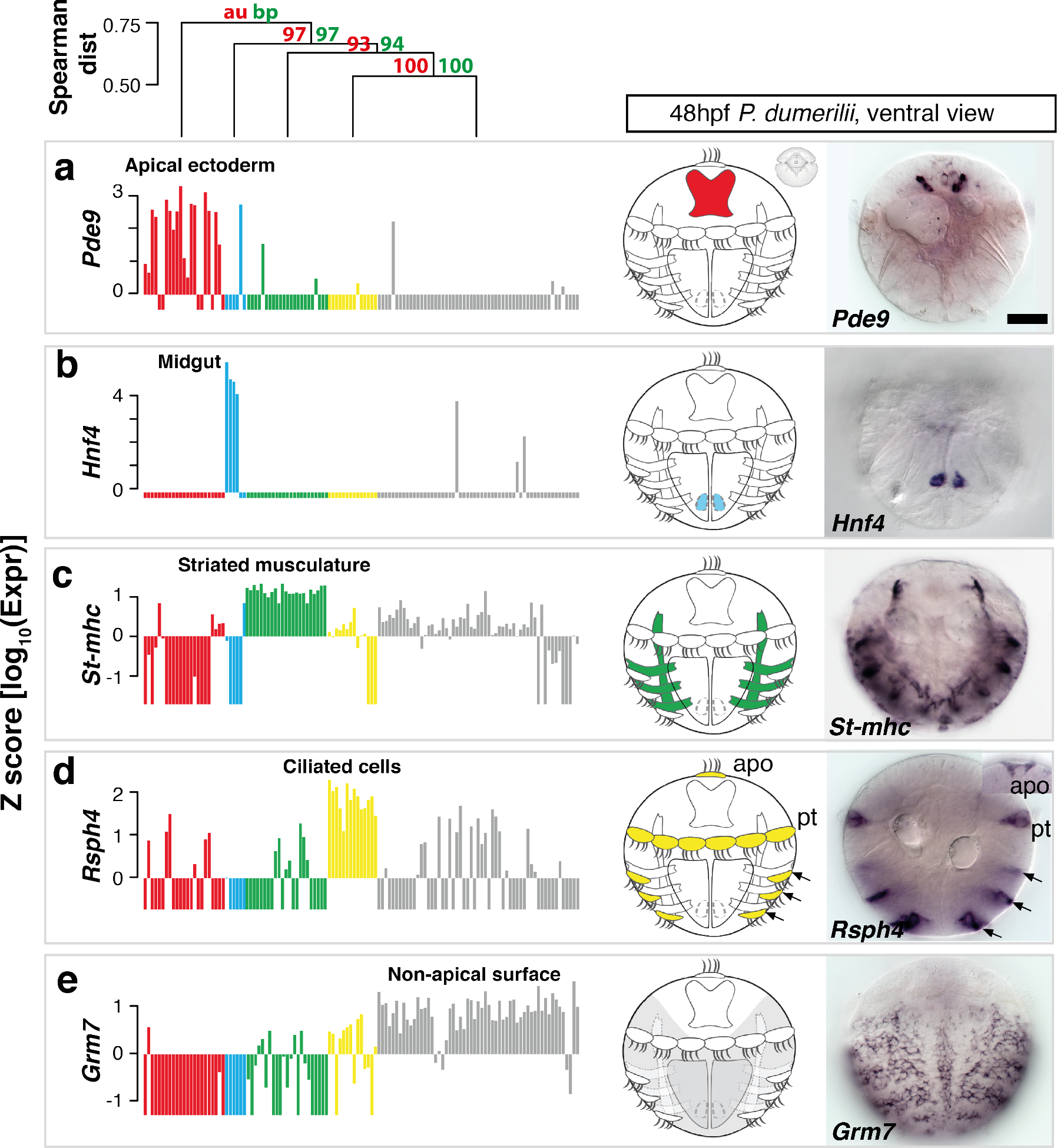
Identification and validation of tissue-specific genes. **On top**, Distance tree showing the hierarchical relationships between the differentiated cell groups. **a-e**, For each identified cell group, the expression of a group-specific marker gene is shown in a bar plot, the respective tissue is shown schematically in the ventral view of *Platynereis* larva, and visualized by WMISH with respective probes: **a**, *Pde9* expression in the apical ectoderm (red); **b**, *Hnf4* expression in the midgut (cyan); **c**, *Stmhc* expression in striated muscle (green); **d**, *Rsph4* expression in ciliated cells (yellow); **e**, *Grm7* expression characterizes the non-apical surface cells (gray). Note that *Pde9, Hnf4, Rsph4* and *Grm7* are novel markers for the respective cell groups. Each ISH pattern was replicated in at least 6 animals. Scale bar, 50 µm. Apo, apical organ; pt, prototroch, Z factor = 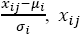: Expression of gene i in cell j, *μ_i_*: Mean expression of gene i, *σ_i_*: Standard deviation of gene i.

Notably, while the sorting of differentiated cells into ciliary bands, larval midgut, and striated musculature matched the subdivision of the larval body into well-known, morphologically distinct tissues, the assignment of ectodermally-derived cells into apical and non-apical populations did not correspond to an obvious morphological boundary. Interestingly, however, the topology of the (unrooted) hierarchical tree (**Fig. 3, Extended Data Fig. 3**) supported the general subdivision of the larval body into apical versus non-apical tissues - suggesting a closer relationship of apical nervous system and midgut on one hand, and of the non-apical tissues (non-apical surface, ciliary bands and striated musculature) on the other.

### The apical nervous system

To further explore the different molecular nature of the five differentiated cell groups, we investigated their distinctive gene sets. **Fig. 4** shows a collection of the most specific and informative regulatory and effector genes specific for each group.

**Figure 4.**
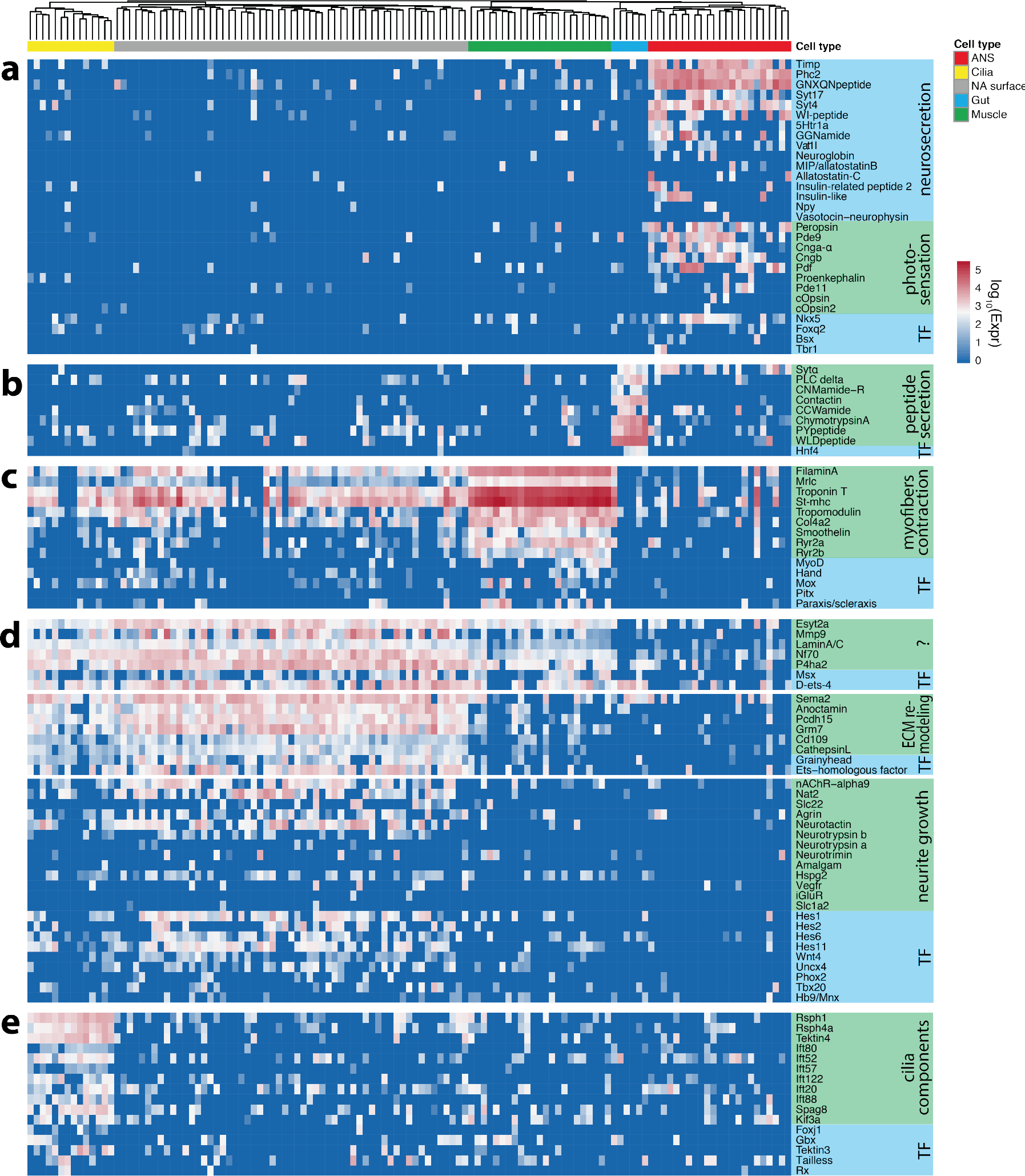
Tissue specific marker genes reflect cellular functions. For each of the differentiated tissues, we show a heatmap of tissue specific gene expression: **a**, expression profile of ANS-specific genes; **b**, expression profile of midgut genes; **c**, expression profile of ciliated cell-specific genes; **d**, expression profile of non-apical surface-specific genes; and **e**, expression profile of myocyte-specific genes. Functionally related groups of genes are highlighted.

Cells of the apical group consistently expressed the neuropeptide-cleaving *Prohormone convertase 2 (Phc2)*^22^ and many neuropeptides, such as the broadly expressed *GNXQNpeptide* or *GGNamide*, two lophotrochozoan neuropeptides^23^, indicating neurosecretory release from these cells. We also noted that cells in the apical group highly expressed non-calcium-binding members of the synaptotagmin family implicated in the generation and fusion of large dense core vesicles for neurosecretion, such as *Synaptotagmin17/B/K (Syt17), Sytα* and *Syt4* (**Fig. 4a-b, Fig. 5, Extended Data Fig. 5**)^24–26^. This is in line with previous reports of neurosecretory cells populating the dorsomedian larval brain^22^, and with ultrastructural observations, which show that cells in the dorsomedian *Platynereis* brain are rich in dense core vesicles while synapses are often sparse or absent ^27^.

**Figure 5.**
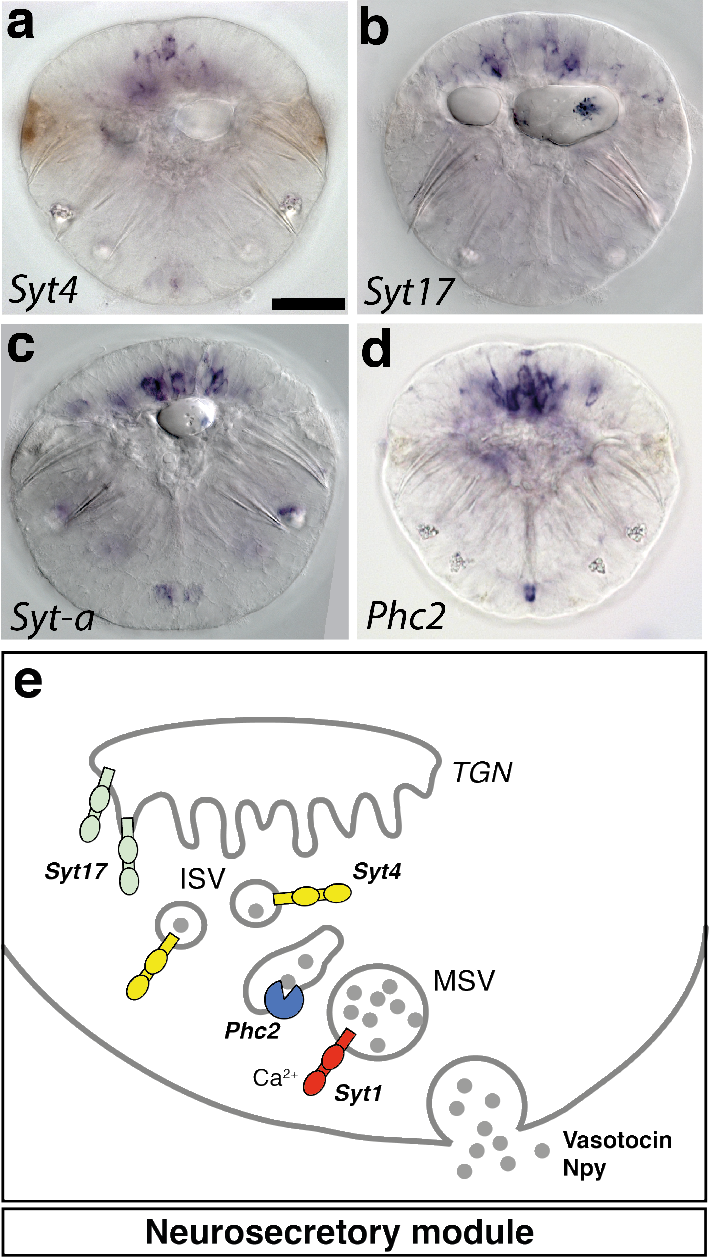
Neurosecretory functional module is specific to ANS cells. a-d, WMISH analysis of *Syt4, Syt17, Sytα* and *Phc2* expression in *Platynereis* larvae at 48hpf. Ventral view. Scale bar, 50 μm **e**, Schematic of the neurosecretion cellular module. Syt17 contains an N-terminal cysteine cluster mediating membrane association with the trans-Golgi network (TGN)^83^. Syt4 is involved in the maturation of secretory vesicles^63^. Sytα functions in vesicle trafficking of specific subclasses of neuropeptides and/or neuromodulators^84^. TGN, trans-golgi network; ISV, immature secretory vesicle, MSV, mature secretory vesicle.

In addition, the bistable photopigment *Peropsin*^28^ and components of cGMP-based signal transduction such as *Pde9* and the CNG channel components *Cngb, Cnga-α* were broadly expressed in the cells of the apical group, indicative of light reception via vertebrate-type phototransduction^29^ (**Fig. 4a**). Specific cells within this group co-expressed effector genes and transcription factors, such as *c-opsin* or *c-opsin2* and *Neuropeptide Y (Npy)* (representing brain ciliary photoreceptors^29^), or *Vasotocin-neurophysin, Proenkephalin* and *c-opsin* (representing photosensory-vasotocinergic cells^22^). Adding to this, some apical cells expressed the neural effector gene *Neuroglobin*, encoding a monomeric globin evolutionarily older than hemoglobin or myoglobin that reversibly binds oxygen and may represent an oxygen reservoir for highly metabolic neurons (**Fig. 4a**)^30^. Other cells expressed *Insulin-like peptide, Pigment dispersing factor (Pdf) or Allatostatin-C*, and subsets of cells expressed the transcription factors *Foxq2, Bsx* and *T-brain* (*Tbr1*). In sum, we conclude that the apical ectodermal cells form part of the photosensory-neurosecretory dorsomedian brain^4,22^ and correspond to the ‘apical nervous system’ (ANS) as defined previously^4,6,27,31^.

### Midgut cells and striated musculature

Cells allocated to the early differentiating midgut specifically expressed the midgut-specific transcription factor *Hnf4* as well as *Chymotrypsin-A*, a digestion enzyme (**Fig. 4b**). In addition, this group was enriched in the expression of signal peptides such as *CCWamide, PYpeptide, WLDpeptide*, indicating that peptidergic neurosecretion is a characteristic feature of this group, similar to the apical nervous system (but using different neuropeptides).

The striated musculature cells were characterized by the relatively higher expression of *St-mhc, Mrlc, Troponin-I* and *Troponin-T*, which together are known to assemble striated myocyte contractile fibres^32^ (**Fig. 4c**), and by the heterogeneous expression of the myocyte-specific terminal selectors *MyoD, Hand, Paraxis, Mox and Pitx*^33^. We used WMISH and co-expression analysis using ProSPr to validate the expression of *MyoD* in a subset of muscles (**Extended Data Fig. 5**). Notably, we detected a low level of the striated myocyte markers in other cell groups (**Fig. 4c**). Given the exceptionally strong expression of these genes in the myocytes, and their up to 100-fold lower expression in the other groups, it is yet unclear whether this finding represents biological or technical noise.

### Non-apical surface cells and ciliary bands

The cells of ectodermally-derived, non-apical surface and the ciliary band cells expressed different, partially overlapping gene sets indicative of mixed epithelial and neural characteristics (**Fig. 4d**). First, we identified a set of genes broadly co-expressed by non-apical surface ectoderm, ciliary bands and - consistently but at lower level – in striated musculature. These genes included *LaminA/C*, a nuclear fibrous protein forming the nuclear lamina on the interior of the nuclear envelope^34^ (**Fig. 4d, Extended Data Fig. 5**), and the related *Neurofilament-70 (Nf70)*, a neural intermediate filament found in Lophotrochozoans (**Fig. 4d**). Interestingly, Lamins and cytoplasmic intermediate filaments evolved via duplication from an ancient, Lamin-like gene (with the intermediate filament precursor losing the nuclear localization sequence and the CaaX motif characteristic for Lamins, and giving rise to intermediate filaments and keratins several times independently^35^. In addition, this group comprised the conserved *Extended Synaptotagmin 2 (Esyt2)*, with an apparent function in membrane lipid composition dynamics, extracellular signal transduction and cell adhesion^36^; and finally, *P4ha2* encodes a component of prolyl 4-hydroxylase, a key enzyme in collagen synthesis (**Fig. 4d**).

Another gene set, shared between non-apical surface cells and ciliary bands, was enriched for extracellular matrix activities. For example, orthologs of *Anoctamin*, a transepithelial chloride channel^37^, *Protocadherin-15 (Pcdh15)*, an atypical cadherin mediating structural integrity of ciliated sensory receptors^38^ (**Extended Data Fig. 5**), and *Semaphorin 2 (Sema2)*, a secreted extracellular guidance molecule involved in axon pathfinding^39^ were co-expressed in these groups (**Fig. 4d**). Most noticeably, however, several genes encoded matrix-modifying proteases such as a serine protease *Plasminogen* related to vertebrate Plasmin^40^ and Hepatocyte Growth Factor (HGF)^41^ (**Extended Data Table 2**); metalloproteinases such as *Matrix Metalloproteinase (Mmp9)*^42^ together with its specific substrate *Cd109*^43^; and the cysteine proteinase *Cathepsin L*^44^ (**Fig. 6b**). Regarding transcription factors, these groups shared the epidermis-specific transcription factor *Grainyhead*, known for its conserved role in maintaining the epidermal barrier^45^, and the *ETS-homologous factor* that likewise specifies epithelial cell type identities^46^ (**Fig. 4d**).

**Figure 6.**
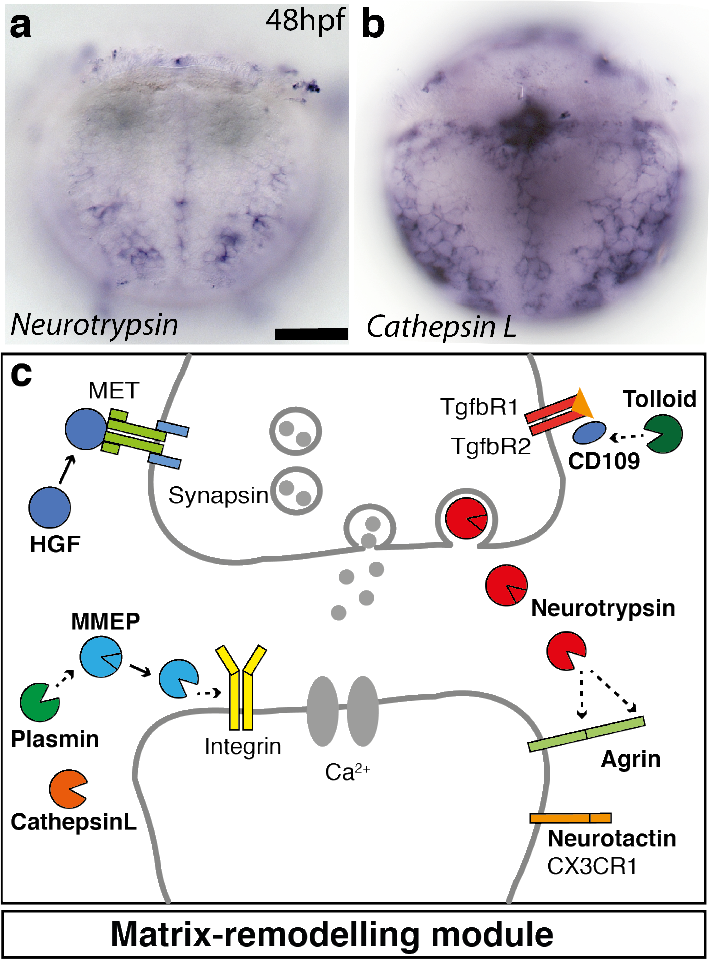
Extracellular matrix remodeling module is specific to non-apical surface cells. **a-b**,WMISH analysis of *Neurotrypsin* and *Cathepsin L* expression in *Platynereis* larvae at 48hpf. Ventral view focused at the surface. Scale bar, 50 μm **c**, Schematic of the synapse formation cellular module. Neurotrypsin cleaves agrin locally at the synaptic cleft, triggering the formation of new synapses^40^. HGF and its receptor MET enhance clustering of synaptic proteins at excitatory synapses^41^. Plasmin cleaves selected synaptic target proteins such as NMDA receptor and matrix metalloproteinases^40^. Several members of the family of matrix-metalloproteinases such as MMP-9 have been in synapse formation and remodeling^42^. *Tolloid-like (Tll)* and its substrate *CD109* have been implicated in the control of TGFb signaling and extracellular matrix synthesis^43^. *Cathepsin L* is likewise implicated in extracellular matrix remodeling and neuronal survival^44^. The genes specifically expressed in non-apical ectodermal cells are marked in bold.

Likewise, the genes most specific for non-apical surface cells abundantly encoded proteins known to specifically interact in extracellular matrix and basal lamina (**Fig. 4d, Fig. 6c**) – many of which with reported roles in nervous system development and plasticity. For example, we identified orthologs of *Neurotactin*, which encodes a cell surface glycoprotein of the serine esterase superfamily^47^, and its specific binding partner *Amalgam* ^48^; both involved in axonal pathfinding in *Drosophila*^47,49^. We also found two orthologs of *Neurotrypsin* (**Fig. 6a**), a matrix-modifying serine protease, together with the extracellular proteoglycan *Agrin*^50^, indicating that the Neurotrypsin-Agrin system is active in these cells^51^ (**Fig. 4d**). The *Hspg2* gene, encoding a conserved proteoglycan related to Agrin, was also present. Besides extracellular matrix components, the non-apical surface cells also expressed *Wnt* ligands such as *Wnt4*^52^, and the bHLH HES transcription factors *Hes1, Hes2, Hes6* and *Hes11*. Homeodomain transcription factors with conserved roles in neuronal specification such as *Uncx4, Tbx20* or *Phox2*^2,53,54^ were expressed in subsets of cells in this group (**Fig. 4d**), indicating neuronal identities.

Finally, we also identified a group of genes most specific for the ciliary bands: *Rsph, Ift, Dynein* and *Kinesin* effector genes, which are constituent parts of motile cilia (**Fig. 4e, Extended Data Fig. 5, Extended Data Table 2**)^55,56^. Notably, all of these genes were also expressed in other groups, albeit in a patchy manner and at lower level. While most of the ciliated cells mapped to the larval trunk, two of the ciliary band cells mapped to the head and co-expressed the transcription factors *Rx* and *Tailless*, which specify head regions and cell types in brain and eye in a broad spectrum of bilaterians ^57^. This suggest that the ciliary band cells may split into distinct subgroups corresponding to the ciliary cells of the head *vs* trunk.

### Distributed nature of neural cells

Interestingly, scRNAseq indicated that pan-neuronal differentiation genes such as *Synaptotagmin1, Rab3, Munc13, Rims, Rimsbp* and *Complexin*^58^ were expressed across all groups of differentiated cells (**Fig. 7b**). This is consistent with observations that neuronal features are not restricted to the nervous system^58^. For *Platynereis*, the ciliated cells of the larval prototroch are reported to show action potentials^59^ and therefore likely to also express neuronal markers; and the *Platynereis* differentiated midgut has recently been show to show autonomous contraction waves at juvenile stages^60^.

**Figure 7.**
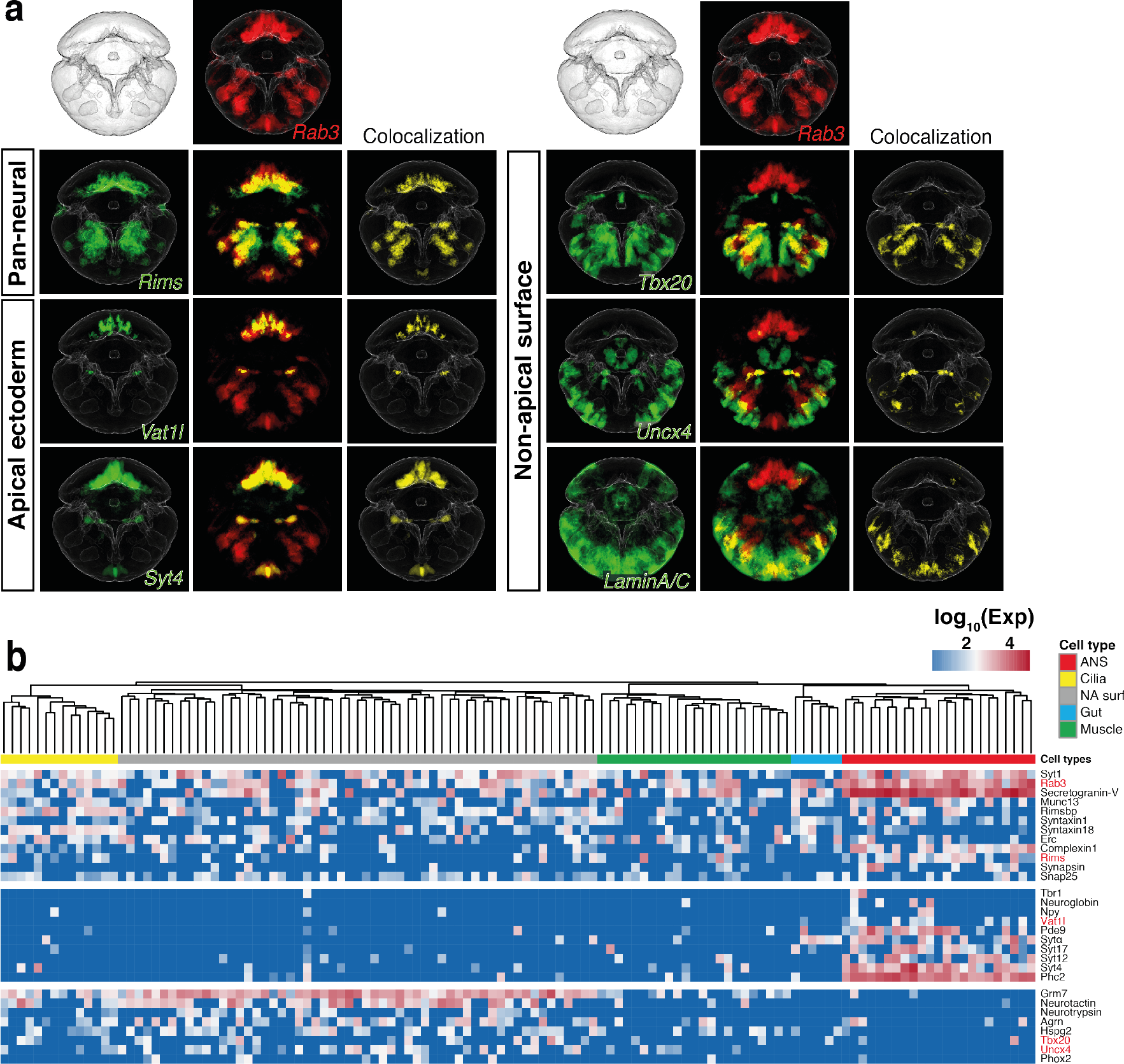
Shared and unique characteristics of the apical and non-apical nervous system. **a**, Averaged gene expression patterns and *in silico* analysis of co-expression with pan-neuronal marker gene *Rab3*, for selected marker genes. **b**, Heatmap showing the expression profile of pan-neuronal markers, as well as neuronal genes specific to apical ectoderm and non-apical surface cells. Red labels, the genes for which ISH is shown in (**a**).

Upon examining the spatial expression pattern of pan-neuronal genes using WMISH we noted that their expression was highest in the apical nervous system and in the ventral portion of the non-apical, ectodermally-derived tissue (**Fig. 7a**), consistent with the known location and extent of the neuroectoderm at this stage^2^. With regard to the apical nervous system, a large fraction of its constituent cells indeed mapped in to the apical domain of *Syt1* expression (**Extended Data Fig. 6a**). Substantiating this, we validated the co-expression of the pan-neuronal marker genes and apical cell group markers by WMISH (**Fig. 7a**).

With regard to the non-apical surface cells, spatial mapping showed that some of those cells also mapped into the *Syt1* expression domain on the ventral body side (**Extended Data Fig. 6b**), supporting their neuronal character. ProSPr analysis likewise confirmed an overlap of non-apical surface markers with the pan-neuronal marker *Rab3* (**Fig. 7a**). Adding to this, we validated the expression of the non-apical group marker *Neurotrypsin* in the ventral neuroectoderm by WMISH (**Fig. 6a**). Together, these data indicate that, at least at this larval stage, ‘neurons’ did not fall into one homogenous group (in contrast to striated myocytes or cells with motile cilia).

## Discussion

For this study we analyzed the transcriptomes of 373 cells randomly collected from dissociated, entire larvae of the marine annelid *Platynereis dumerilii*. Among these we identified 123 differentiated cells, chosen for further analysis and comparison. Previous studies had captured cells from specific body parts and tissues only (such as the mouse forebrain^9,10^); our data is unique as it allows single-cell level comparison of differentiated cells across the entire body. We were able to sort all differentiated cells into five major groups, based on similarities and differences in gene expression profiles. Spatial mapping into the 48 hpf ProSPr cellular-resolution expression atlas, in combination with wholemount in situ hybridisation of marker genes, identified these groups as apical nervous system, larval midgut, ciliary bands, non-apical surface cells, and striated musculature.

What is the nature of these groups? Interestingly, within each of the five groups, there existed considerable heterogeneity in the expression of transcription factors and effector genes (as evident from **Fig. 4**), suggesting the presence of several cell types within each group. For example, cells of the apical nervous system differ with regard to opsin expression, neuropeptide content, and signaling components – indicating that each, or almost each of the cells might represent a separate type. Therefore, at the current level of analysis the five groups of differentiated cells rather appear to represent some kind of molecularly defined larval tissues – uniting cells with similar molecular properties shared beyond the cell type level. We believe that the number of sequenced cells would have to be increased by at least one order of magnitude to resolve individual cell types within these groups.

Comparing the cell groups further, we noted an unexpected divide between ectodermally-derived cells in the most apical part of the larva (apical nervous system), *versus* the non-apical surface cells and ciliary bands. Cells of the apical nervous system shared the expression of two major sets of modules, enabling the synthesis and secretion of neuropeptides, and mediating the detection of ambient light for the circadian control of body physiology. Most of the specifically expressed effector genes detected for these modules, encoding for example prohormone convertase, specific neuropeptides, components of the phototransductory cascade, and ciliary opsins, have previously been implicated in apical nervous system function^4,22^.

In contrast, the genes characteristic for the non-apical surface cells revealed mixed epithelial and neural properties. One broadly expressed set of genes – found in non-apical surface, ciliary band and striated muscle cells – encoded proteins relating to basic properties of the surface epidermis – such as stabilizing intermediate filaments (related to vertebrate keratins), and an enzyme mediating collagen synthesis into the extracellular matrix. Another set of genes – shared between non-apical surface and ciliary band cells, but absent from myocytes – included matrix-modifying proteinases, many of which implicated in tissue remodeling, axonogenesis and synapse formation in insects and vertebrates^40^. And yet another set – identified as most specific for the non-apical surface cells proper – included genes encoding neural properties, such as *Neurotactin/Amalgam, Neurotrimin*, and *Semaphorin* involved in axon guidance and *Neurotrypsin/Agrin* with reported roles in synaptogenesis^51,50^. This would indicate that the non-apical surface cells utilize specific molecular toolkits to modify extracellular matrix, guide outgrowing axons, and establish synapses – that are not employed in the apical nervous system.

The specific and broad expression of the nicotinic acetylcholine receptor alpha-9 in non-apical surface cells is also noteworthy. In vertebrates, this unconventional receptor plays restricted roles in the mechanosensory hair cells, where it is involved in the reception of efferent signals^61^, and in epidermal keratinocytes, where it triggers epithelialization via local, non-neuronal acetylcholine cytotransmission^62^.

What is the significance of the fundamental divide between apical and non-apical cells in the *Platynereis* larval surface ectoderm? While some of the genes that are differentially expressed between apical and non-apical cell groups relate to functions carried out by fully differentiated cells (e.g., neurosecretion, phototransduction, cholinergic cytotransmission), others indicate earlier stages of differentiation (e.g., proteolytic cleavage for extracellular matrix modification, and axonal guidance). Thus, and given that our analysis only covers one single stage of larval development, we cannot rule out that some of the differences between apical and non-apical cells are due to differences in developmental timing; more extensive studies including additional stages will be required to solve this issue. However, preliminary WMISH of ‘apical’ and ‘non-apical’ marker genes already indicates that expression differences between apical and non-apical tissues are also observed at earlier developmental and later life-cycle stages (**Extended Data Fig. 7**). Developmental timing is therefore unlikely to fully account for the apical/non-apical differences.

Another possible explanation for the difference between apical and non-apical cells is evolutionary divergence. This notion assumes that an ancient subdivision of the animal body into functionally and structurally different parts occurred early in animal evolution. These segregating parts would have started to differ in epithelial properties, extracellular matrix remodeling capacities and cell-cell communication strategy. To validate and substantiate this notion, similar datasets from both, closely and remotely related species will be required; they should show similar overall molecular tissue subdivisions if these were evolutionarily conserved.

As a starting point, and in line with the second option, the restricted activity of apical nervous system marker genes in the vertebrate forebrain indicates evolutionary conservation of the apical nervous system and of its specific properties^6^. For example, in vertebrates the neurosecretory-specific *Syt4* is specifically active in the neuroendocrine hypothalamus, where it regulates release of the neuropeptide Oxytocin^63^. *Syt17*, initially isolated from the vasopressin-secreting supraoptic and paraventricular nuclei, also shows strong expression in the hypothalamus^64^, as does *Neuroglobin*^65–67^. The T-box transcription factor *T-brain* is specifically expressed in the developing para- and periventricular and supraoptic hypothalamic nuclei^68^; and finally, expression of *ciliary opsins* and other genes of the ciliary photosensory cascade in the vertebrate retina and pineal indicates evolutionary conservation of apical nervous system components^29^. These similarities add to previously reported conserved features shared between vertebrate hypothalamus and annelid apical nervous system, involving specifying transcription factors^69^, a neurosecretory centre releasing conserved neuropeptides and hormones^22,27^, and non-visual ciliary photoreceptors^29^ active in melatonin synthesis and the control of circadian behavior^4^. Our transcriptomics data thus support the evolutionary conservation of the apical nervous system as a distinct part of the bilaterian brain, specialized on the perception of ambient light for the control of body physiology via the release of neuropeptides and hormones. They are furthermore consistent with the recent ‘chimeric brain’ model, which stipulates that the apical nervous system evolved as an independent center of neuronal condensation in the course of bilaterian brain evolution^6^.

## METHODS

### Cell capture and sequencing

*Platynereis* larvae were cultured, the cells were dissociated, and single-cell cDNA synthesis was performed using the Fluidigm C1 Single-Cell Auto Prep System, and the sequencing libraries were prepared as previously described^5^. We used Fluidigm IFCs for mRNA-seq with capture sites for 5-10 µm and 10-17 µm to collect the cells, allowing the capture of a full range of cell sizes in 48hf stage of *Platynereis*. In total, 9 IFC-s, each representing an independent collection of cells derived from a different batch of animals, were processed for this study. The information about the samples distribution on chips, the chip sizes, cell number per sample and the pooling of samples on sequencing lanes described in Extended Data Table 3. 100 bp paired-end sequences were generated using the Illumina HiSeq2000 platform. For one library (CN62), 75 bp paired-end sequences were generated using the Illumina NextSeq500 platform. Libraries were sequenced to an average depth of 2.1 million reads.

### Data availability

The raw sequence data for the cells analysed in this study are available from ArrayExpress, accession numbers E-MTAB-2865 (see **Extended Data Table 5** for matching the cell ID-s between the E-MTAB-2865 and this study) and E-MTAB-XXXX (the accession number will be updated as the dataset is released). The R code used for analysis is available at: https://github.com/MarioniLab/Platynereis2017

### Read alignment

FastQC^70^ was used for the quality trimming of the raw sequencing reads using default settings. For read mapping, the quality-filtered sequencing reads were mapped, using bowtie2, against a *Platynereis* reference transcriptome combined from the two previously published assemblies^5,23^. Briefly, the transcriptome assemblies were concatenated and contigs showing more than 94% identities were removed using CD-HIT^71^, leaving 44977 transcripts. In thus generated reference, each gene should be represented by one transcript, reducing the problem of multiple mapping of the reads. Expression counts for each transcript were obtained using HTSeq^72^ while only considering uniquely mapped reads.

### Gene annotation

To annotate genes of the two transcriptome assemblies^5,23^, reciprocal BLAST comparison of genes sequences against the Uniref90 protein database (Arendt assembly) or against Swissprot (Jékely assembly) was performed. For each transcript, the BLAST hit with highest E-value was selected for annotation.

### Quality control

Low-quality cells were removed from the dataset based on the following filtering criteria. Visual inspection of capture sites on the IFC revealed empty wells, wells with multiple cells or debris captured. Only wells with positive cell capture were further processed for sequencing. For downstream analysis, libraries marked as single cells were chosen. Cells with less than 1000 reads mapping to ERCC spike-ins, less than 100,000 mapped transcriptomic reads or less than 60% of mapped reads being allocated to the transcriptome were removed. Additionally, cells that express unusually small number of genes (< 2000) were removed. Prior to normalization, genes with more than 1,000,000 reads or less than 10 reads across all cells were removed.

### Normalization

The BASiCS package^73^ was used to normalize read counts by incorporating ERCC spike-ins for technical noise estimation. Specific ERCC spike-ins were removed if not detected in the dataset. Posterior estimates for model parameters were computed by Markov chain Monte Carlo (MCMC) simulation with 40,000 iterations.

The *BASiCS_DenoisedCounts* function was utilized to compute normalized read counts. Running the MCMC independently on the different batches confirmed similar small levels of technical noise in the data (**Extended Data Fig. 1h**),

### Spatial mapping

Spatial mapping of the sequenced single-cell transcriptomes onto the *Platynereis* reference atlas was performed as described previously^5^. The *Platynereis* reference atlas is provided as **Extended Data Table 4** and the spatial mapping results for all the sequenced cells are provided in **Extended Data Table 5.** We defined functional regions within the embryo based on spatial expression patterns of known marker genes. The region in which *Phc2* is expressed comprises the apical nervous system. The muscle region of the animal is defined by the expression of *St-mhc. Foxj* expression defines the ciliated cells and *Hnf4* expression the midgut region. The trunk ectoderm can be described by *Uncx4* expression. To assign mapped cells to particular regions, we identified the voxel cluster with highest mapping confidence for each cell. For visualizing the mapping of individual cells, the centroids of all voxel clusters were plotted. To compute the overlap of mapped cells to spatial gene expression patterns, we focused on the cluster with highest mapping confidence for each cell. If more than 50% of these voxels also show a particular gene expression (e.g. *Syt1*), we consider the cell as falling into this gene region.

### Detecting highly variable genes

The *BASiCS_DetectHVG* function in BASiCS allows the detection of highly variable genes by incorporating spike-ins to estimate expected technical variability. Testing was done after MCMC simulation with a variance threshold of 0.98 and an evidence threshold of 0.7.

### Clustering

To detect clusters of cells based on the expression of highly variable genes, we used a sparse K-means clustering approach taken from the *sparcl* package in R^74,75^. We clustered the data using K=2,…,10 and chose K=7 since only small subpopulations emerged at K=8,…,10. An elbow plot showing the averaged within-cluster sum of squares is non-informative for the choice of K. *AWSS* = (Σ_*k*_Σ_*i*_Σ_*j*_(*x_ijk_* – *μ_ik_*)^2^)/*m*, where *x_ijk_* is the log_10_-transformed transcript count of gene i in cell j and cluster k, *μ_i_* the log_10_-transformed mean expression of gene i in cluster k and *m* the number of clusters. The parameters were tuned with 2 < wbounds < 100 using 15 permutations. Genes with weights > 0.03 define the cluster-characteristic features of the data.

### Removing cell doublets

Possible cell doublets were removed after clustering and marker gene detection. The percentage of group specific marker gene expression (# marker genes expressed/# all genes) was calculated for each cell. Cells with unusually high expression of markers non-specific for their group were removed from the dataset.

### Tree building

To investigate the transcriptional similarity between cell types, we hierarchically clustered cell types before and after averaging gene expression across all cells within each group. The averaging strategy increases stability of the hierarchical clustering algorithm by averaging out dropout events in single cell data. Distances were calculated based on the Spearman dissimilarity: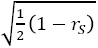^76^. To compute bootstrapped p-values, we used the *pvclust* package in R^77^. Hierarchical clustering was performed on the log_10-_transformed expression counts before and after averaging within each group using the spearman dissimilarity as distance, complete linkage clustering and 1000 bootstrap iterations to evaluate cluster stability.

### Differential expression

To identify cell group specific enriched marker genes, we used two approaches. First, we used scde package^78^ to compare each of the identified cell group against all the remaining samples in order to identify novel group specific genes from the unannotated portion of the Platynereis transcriptome. These genes were then annotated, added to the reference, and the clustering was repeated. Second, differential expression analysis was performed using the *findMarkers* function from the *scran* package^62^. This approach uses *limma* ^79^ on the log_2_-transformed, normalized counts. Group specific marker genes were defined as differentially expressed genes with a FDR < 0.1 and log_2_FC > 0 for all pairwise comparisons. Initially, differential expression was tested between all 7 detected groups. Group 6 and group 7 consistently grouped together when hierarchical clustering (default options from the *hclust* package) was performed on the log_2_ fold changes in expression between one group and all others (**Extended Data Fig. 2**). We therefore merged group 6 and group 7 and tested differential expression between the now largest group and all other groups (**Extended Data Table 1**). In the next step, we tested differential expression pairwise between all differentiated cell groups (**Extended Data Table 2**).

### *In situ* hybridization

For *in situ* hybridization (ISH), we collected 48 hpf or 72 hpf *Platynereis* larvae. Animals were fixed overnight in 4% PFA/1.5xPBS and ISH was performed as described previously^80^.

### Cloning of *Platynereis* genes and ISH probe synthesis

For the cloning of *P. dumerilii* cDNA sequences, wild-type *Platynereis* RNA was isolated by Trizol/phenol/chloroform extraction method, reverse transcribed using SuperScriptIII reverse transcriptase (Life Technologies, cat. # 18080044), and amplified by PCR using TaKaRa ExTaq DNA polymerase (Clontech, cat. # RR001A) and respective gene specific primers. The resulting PCR fragments were cloned into pCRII-TOPO vector (Life Technologies, cat # K4610-20). The cloned gene sequences and the gene specific primer sequences are available upon request.

For the synthesis of ISH probes, cDNA plasmids were linearized and antisense RNA probes were transcribed using SP6 or T7 RNA polymerase (Roche, cat. #11487671001 and 10881775001, respectively) and DIG RNA-labeling mix (Roche, cat. #11277073910) or Fluorescein RNA labeling mix (Roche, cat. # 11685619910).

### Microscopy and image processing

For imaging of WMISH samples for ProSPr, samples were mounted in 97% 2,2′-thiodiethanol (Sigma, cat. # 166782) and imaged on a Leica TCS SP8 confocal microscope, using a combination of fluorescence and reflection microscopy^81^. The colocalization analyses and image post-processing was performed using Fiji^82^ software. The colorimetric WMISH samples were imaged on Zeiss AXIO Imager M1 microscope. The figure panels were compiled using Adobe Illustrator and Adobe Photoshop software.

## Acknowledgements

We thank Catalina Vallejos for comments on the implementation of BASiCS and all members of the Marioni and Arendt labs for feedback on the project and manuscript.

## Author contributions

KA and NE planned and conducted experiments, interpreted the data, designed the study and wrote the manuscript. HMV, PB and PC conducted experiments. TB interpreted data. VB, JCM and DA funded the study. JCM and DA designed the study and wrote the manuscript.

## Author information

The authors declare no competing interest.

